# SNP-Slice Resolves Mixed Infections: Simultaneously Unveiling Strain Haplotypes and Linking Them to Hosts

**DOI:** 10.1101/2023.07.29.551098

**Authors:** Nianqiao P. Ju, Jiawei Liu, Qixin He

## Abstract

Multi-strain infection is a common yet under-investigated phenomenon of many pathogens. Currently, biologists analyzing SNP information have to discard mixed infection samples, because existing downstream analyses require monogenomic inputs. Such a protocol impedes our understanding of the underlying genetic diversity, co-infection patterns, and genomic relatedness of pathogens. A reliable tool to learn and resolve the SNP haplotypes from polygenomic data is an urgent need in molecular epidemiology. In this work, we develop a slice sampling Markov Chain Monte Carlo algorithm, named SNP-Slice, to learn not only the SNP haplotypes of all strains in the populations but also which strains infect which hosts. Our method reconstructs SNP haplotypes and individual heterozygosities accurately without reference panels and outperforms the state of art methods at estimating the multiplicity of infections and allele frequencies. Thus, SNP-Slice introduces a novel approach to address polygenomic data and opens a new avenue for resolving complex infection patterns in molecular surveillance. We illustrate the performance of SNP-Slice on empirical malaria and HIV datasets and provide recommendations for the practical use of the method.

## 1 Introduction

Multi-strain infection is a common yet under-investigated phenomenon of many viral, bacterial, fungal, and protozoan diseases [1]. For example, 30-80% of individuals in endemic malaria transmission regions often carry multiple, genetically-distinct strains of the same pathogen species [2, 3]. The presence of multiple strains in individuals often results in poor treatment outcomes [4] or increased virulence from within-host competition [5]. Current detailed analyses of population-level parasite diversity, relatedness, and regions subject to selection require strictly monogenomic infection inputs. Therefore, isolates with a high complexity of infection are often excluded from these analyses, which amounts to the majority of samples from high-endemic regions [6, 7]. In addition to the unfortunately wasted resources, discarding mixed infection samples leads to selection biases in data analysis. More importantly, it remains difficult to understand the parasite diversity patterns in high-endemic regions, where the knowledge is most needed. Therefore, a reliable tool to learn and resolve the SNP haplotypes from polygenomic data is an urgent need in molecular epidemiology.

Earlier algorithmic efforts on resolving haplotypes started with phasing diploid genotypes in human genetics [8, 9], and later were extended to pooled samples [10–12]. Nevertheless, these methods do not apply to inferring many co-infected haplotypes in microorganisms because the underlying genetic mechanisms between humans and microbes, such as coalescence and recombination patterns, are very different, and the methods do not scale up to haplotype reconstruction for more than 25 SNPs. With the advancement of sequencing technologies, many algorithms were designed to utilize physical linkage of SNPS on reads for microbial metagenomic studies [13–15]. They often rely on deep sequencing coverage per strain (> 50×) or existing strain reference libraries [16]. However, field samples with mixed infections usually contain low concentrations of parasite DNA [17], and correspond poorly to reference libraries. Recent reference-free methods are limited to 2-4 strains per infection [18, 19] or limited to haplotypes of a few SNPs [20]. In addition, most of these algorithms focus on resolving mixed infections within one host [13–15, 18], without utilizing information from population-level haplotypes and allele frequencies. Another class of algorithms focused on learning the number of strains infecting each host, known as the multiplicity of infection (MOI), and allele frequencies at the population level, without actually resolving the constituent haplotypes [21–24]. Although these methods provide an estimation of important epidemiological quantities, it is unsatisfying because most of the fine-scale molecular genetic analyses require haplotype information. Thus, it is imperative to develop methods with a focus on population-level inference of haplotypes, without heavy parametrization, to ensure a wide application to different diseases.

In this work, we develop SNP-Slice, a family of slice sampling Markov Chain Monte Carlo algorithms, to infer SNP haplotypes infecting the population as well as whether they appear in each host. In other words, we aim to fully resolve all multi-strain infection information in terms of the haplotype dictionary (feature extraction) and host-strain association (feature allocation). In addition, many epidemiological characteristics, such as MOI, allele frequency, heterozygosity, and *F*_*WS*_ can be derived from our inference results (Section 4.5). In the SNP-Slice (SS) framework, we use the Indian Buffet Process (IBP) prior [25, 26] to describe host-strain associations (Section 4.3), 2 which imposes a clustering structure among the hosts and strains, and allows each host to be infected with any combination of the (potentially infinitely many) strains. IBP priors have been applied successfully in a large variety of research fields including imaging segmentation [26], natural language processing [27], psychology [28], and more biologically relevant ones such as biomarker discovery [29], studying protein interactions [30] and gene expressions [31]. In IBP, a single concentration parameter *α* describes our prior belief about the host-strain clustering structure. Adopting the IBP prior, the complexity of SNP-Slice is data-adaptive, since the size of the inferred haplotype dictionary can grow as we observe more data. This makes the SS method simpler to tune than competing methods that require more model parameters. Moreover, we propose and examine three novel observational models (Bin, Pois, and NegBin) that tailor to SNP read counts data, which accommodate heterogeneous sampling efforts across SNP sites (Section 2.4) and unbalanced within-host strain frequencies (Section S1.2). We also introduce an SS-Cat method to study SNP categorical data that indicates the heterozygosity status at each site. In sum, the SNP-Slice framework includes four algorithms: SS-Bin, SS-Pois, SS-NegBin, and SS-Cat. In terms of computational contributions, SNP-Slice efficiently solves the challenge of high-dimensional Bayesian inference and yields accurate reconstruction of the haplotype dictionary as well as their presence in hosts. Extensive experiments demonstrate the effectiveness of the proposed methods on simulated malaria mixed infection datasets. Based on our results, we recommend that analysts use SNP read count data when available and that SS-NegBin or SS-Cat be used to analyze field data (SS-NegBin for high transmission regions and SS-Cat for low transmission regions). From an evolutionary biology perspective, the SNP-Slice framework will help recover unexplored information from molecular surveillance of disease transmission, evaluation of transmission control, and detailed monitoring of disease progression within hosts. We demonstrate its performance on HIV-1 time series data (Section 2.6) and malaria field datasets (Section 2.5) and its ability to discover new biological insights from resolved haplotype information.

## 2 Results

### 2.1 SNP-Slice outperforms existing methods at MOI and allele frequencies estimations across simulation scenarios and data types

The SNP-Slice framework simultaneously learns the haplotype dictionary (*D*) and its host-strain assignment (*A*). We performed comprehensive benchmarks on realistic synthetic datasets across scenarios of varying transmission intensity that span low to high transmission zones of malaria, using an agent-based stochastic simulator [32] (Section S2.3).

First, we compared SNP-Slice’s performance to the state-of-the-art MOI estimation method The Real McCOIL (hereafter RM) [22]. Examining the error of the estimated MOI against the true MOI 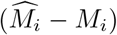, we find SNP-Slice to be more accurate than RM, both on the population level (Table 1) and stratified by true MOI values (Figure 1). In particular, SNP-Slice achieves exact MOI recovery 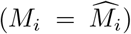 for at least 60% of the individuals, across simulated infection scenarios; and our estimations are within ±1 from the truth for more than 75% of the individuals (Table 1). Table 1 also reports the bias and root-mean-squared error (RMSE)of MOI estimations. When the transmission rate is low (Scenario 1), inference algorithms using heterozygosity status input (SS-Cat and RM-Cat) yield better MOI estimation than algorithms using SNP count input, and 99.8% of MOI estimates from SS-Cat are within ±1 of the truth. As the transmission rate increases (Scenario 2&3), RM-Cat and SS-Cat begin to underestimate MOI. Under the highest transmission intensity, while RM-Prop makes an average error of -1.2, SS-Pois and SS-NegBin can control estimation error below 0.4 in absolute value. In Figure 1, where we group hosts based on true MOI values in Scenario 2, all methods of SS eventually underestimate MOI as the true MOI increases above 5. RM-Prop has a higher negative bias in estimation when compared to SS-Pois and SS-NegBin. For both RM-Prop and RM-Cat, the inferred MOI becomes increasingly variable as the true MOI increases.

**Table 1.** Summary of MOI estimations: mean inferred MOI 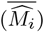, percentage of individuals with exact MOI recovery, percentage of individuals with MOI error *< ±*1, bias and RMSE under different malaria transmission intensities.

| Method | $\widehat{M}_i$ | % of $M_i = \hat{M}_i$ | % of $\vert M_i - \hat{M}_i\vert \leq 1$ | Bias | RMSE |
| --- | --- | --- | --- | --- | --- |
| Scenario 1, average MOI=1.27, $\alpha=1.5$ | | | | | |
| RM-Prop | 1.37 | 91.4 | 97.2 | 0.10 | 0.43 |
| SS-Bin | 1.54 | 87.2 | 91.5 | 0.27 | 0.84 |
| SS-Pois | 1.47 | 89.8 | 93.4 | 0.20 | 0.69 |
| SS-NegBin | 1.40 | 92.2 | 96.0 | 0.13 | 0.53 |
| RM-Cat | 1.35 | 92.1 | 98.6 | 0.08 | 0.36 |
| SS-Cat | **1.27** | **97.4** | **99.8** | **-0.004** | **0.20** |
| Scenario 2, average MOI=2.58, $\alpha=3.0$ | | | | | |
| RM-Prop | 1.94 | 70.0 | 80.0 | -0.64 | 1.73 |
| SS-Bin | 3.46 | 66.2 | 73.9 | 0.88 | 1.88 |
| SS-Pois | 2.90 | 70.3 | 82.3 | 0.32 | 1.26 |
| SS-NegBin | **2.68** | 74.7 | **86.2** | **0.10** | **1.10** |
| RM-Cat | 2.27 | 68.4 | 80.9 | -0.31 | 1.58 |
| SS-Cat | 2.00 | **76.2** | 83.7 | -0.58 | 1.48 |
| Scenario 3, average MOI=3.90, $\alpha=4.0$ | | | | | |
| RM-Prop | 2.69 | 58.0 | 72.0 | -1.20 | 2.90 |
| SS-Bin | 4.84 | 59.1 | 72.6 | 0.94 | 2.08 |
| SS-Pois | **3.70** | 60.3 | **75.8** | **-0.20** | **1.89** |
| SS-NegBin | 3.49 | 61.5 | **75.8** | -0.41 | 2.04 |
| RM-Cat | 3.42 | 57.9 | 75.1 | -0.48 | 2.31 |
| SS-Cat | 2.58 | **64.3** | 73.3 | -1.32 | 2.83 |

**Fig. 1.**
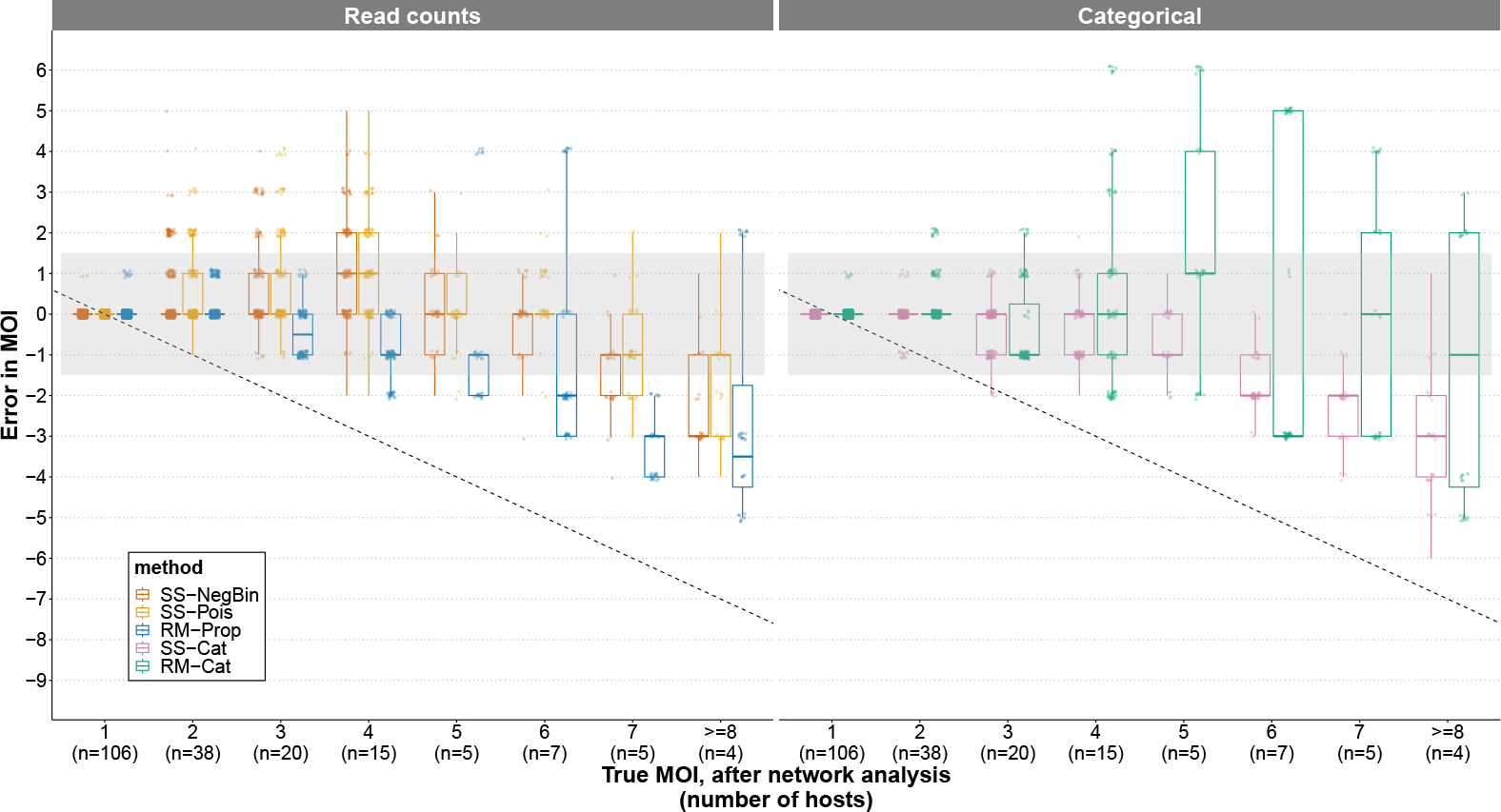
Comparison of SNP-Slice’s MOI estimation performance to that of The Real McCOIL, stratified by true MOI. Boxplots of error in MOI estimation 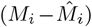 in Scenario 2 are shown. The shaded region indicates an estimation error within ±1. These results summarize 128 independent SNP-Slice repetitions after removing identical duplicate strains for each individual, and network analysis has been performed on true and inferred MOI values by merging strains that share identical SNP type at ≥ 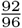 of total sites with others. Each SNP-Slice chain length is set to 10,000 iterations if the input data is read counts (SS-NegBin and SS-Pois), and 5,000 iterations if the input data type is categorical (SS-Cat). The RM results are also based on 128 repeats with 10,000 iterations each. More details about the parameters for running the algorithm are listed in Section S2.5. For visual clarity, a random subset of 5% points in each case is shown on top of the boxplots.

SNP-Slice also outperformed RM at estimating allele frequencies (AF; Section 4.5). We reported the bias, RMSE, and relative error [RE, the ratio of residual sum of squares (RSS) to total sum of squares (TSS)] of their allele frequency estimations in Table 2. While all methods appear to underestimate AF (see negative biases in Table 2), the SS-based methods have smaller AF estimation errors than RM-based methods across all transmission intensity levels. Count-based methods are more accurate than RM-Cat and SS-Cat, which uses categorical input. Notably, SS-Cat achieves higher estimation accuracy than RM-Prop, whose input, SNP read counts, is actually more informative than that of SS-Cat. This demonstrates SS’s ability to effectively extract information and interpret mixed infection data.

**Table 2.** Bias, RMSE, and relative error of allele frequency estimations. RE is calculated with RSS/TSS and reported in percentage.

| Method | Scenario 1 |  |  | Scenario 2 |  |  | Scenario 3 |  |  |
| --- | --- | --- | --- | --- | --- | --- | --- | --- | --- |
|  | Bias | RMSE | RE | Bias | RMSE | RE | Bias | RMSE | RE |
| RM-Prop | -0.012 | 0.019 | 2.31% | -0.047 | 0.092 | 62.6% | -0.044 | 0.090 | 75.5% |
| SS-Bin | -0.008 | 0.020 | 2.79% | **-0.009** | **0.021** | **3.18%** | **-0.019** | **0.035** | **11.5%** |
| SS-Pois | -0.005 | 0.016 | 1.70% | -0.014 | 0.031 | 7.20% | -0.022 | 0.040 | 14.9% |
| SS-NegBin | **-0.004** | **0.013** | **1.07%** | -0.015 | 0.029 | 6.45% | -0.021 | 0.038 | 13.7% |
| RM-Cat | -0.012 | 0.019 | 2.41% | -0.061 | 0.107 | 85.5% | -0.063 | 0.110 | 114% |
| SS-Cat | -0.006 | 0.017 | 1.94% | -0.041 | 0.092 | 63.9% | -0.036 | 0.088 | 72.3% |

### 2.2 SNP-Slice accurately recovers heterozygosity of individuals

We investigated SNP-Slice’s ability to learn the individual heterozygosity per locus, which reflects the genetic diversity of strains within a host (Section 4.5). We highlight that only inference methods that recover both strain dictionary and strain allocations are able to provide heterozygosity estimations. As a result, competing methods such as RM cannot perform such an important task. As shown in Table 3, SNP-Slice recovers individual heterozygosities with high precision. Although all methods slightly over-estimate heterozygosity, the highest estimation bias across all malaria transmission intensities is only 0.008 (RE=4.67%) with the count-based methods, with increasing accuracy as transmission intensity decreases.

**Table 3.** Bias, RMSE and RE of individual heterozygosity estimations.

| Method | Scenario 1 |  |  | Scenario 2 |  |  | Scenario 3 |  |  |
| --- | --- | --- | --- | --- | --- | --- | --- | --- | --- |
|  | Bias | RMSE | RE | Bias | RMSE | RE | Bias | RMSE | RE |
| SS-Bin | **0.0003** | **0.0034** | **0.145%** | **0.003** | **0.009** | **0.805%** | 0.008 | 0.025 | 4.667% |
| SS-Pois | 0.0008 | 0.0045 | 0.254% | 0.003 | 0.010 | 0.946% | **0.005** | **0.023** | **3.956%** |
| SS-NegBin | 0.0007 | 0.0049 | 0.303% | 0.003 | 0.012 | 1.285% | 0.005 | 0.025 | 4.919% |
| SS-Cat | 0.0011 | 0.0119 | 1.754% | 0.008 | 0.031 | 9.292% | 0.021 | 0.052 | 20.23% |

### 2.3 SNP-Slice accurately recovers the strain dictionary

While accurate heterozygosity estimations already imply an accurate recovery of the haplotype dictionary, we further studied the quality of the inferred haplotypes learned by comparing them with the underlying true haplotypes. First, we define the dissimilarity between two haplotypes to be the proportion of sites with different alleles: Two identical haplotypes have a dissimilarity of 0 and two opposite haplotypes have a dissimilarity of 1. Naturally, the similarity between two haplotypes is one minus their dissimilarity. Second, we investigated how many of the true haplotypes are recovered well by SNP-Slice. We define the recovery score of each true strain to be the largest similarity between this true strain *D*_*k*_ and any inferred strain 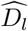 in the dictionary learned by SNP-Slice:

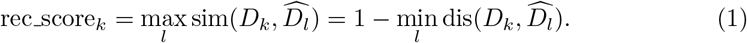

In Table 4, we report the percentage of true strains with a recovery score higher than various cutoff values for Scenario 2, which is our mid-intensity setting. We observed that count-based methods have similar performance and recovering haplotypes becomes more challenging as the transmission intensity increases. Under low transmission rate, more than ≈ 78% of the strains can be exactly recovered by SS-NegBin, and for ≈99% of the true haplotypes we can find an inferred haplotype that is more than 80% similar to it. Even in the highest transmission intensity situation, more than ≈ 72% of the true haplotypes are recovered at the 80% cutoff value.

**Table 4.**
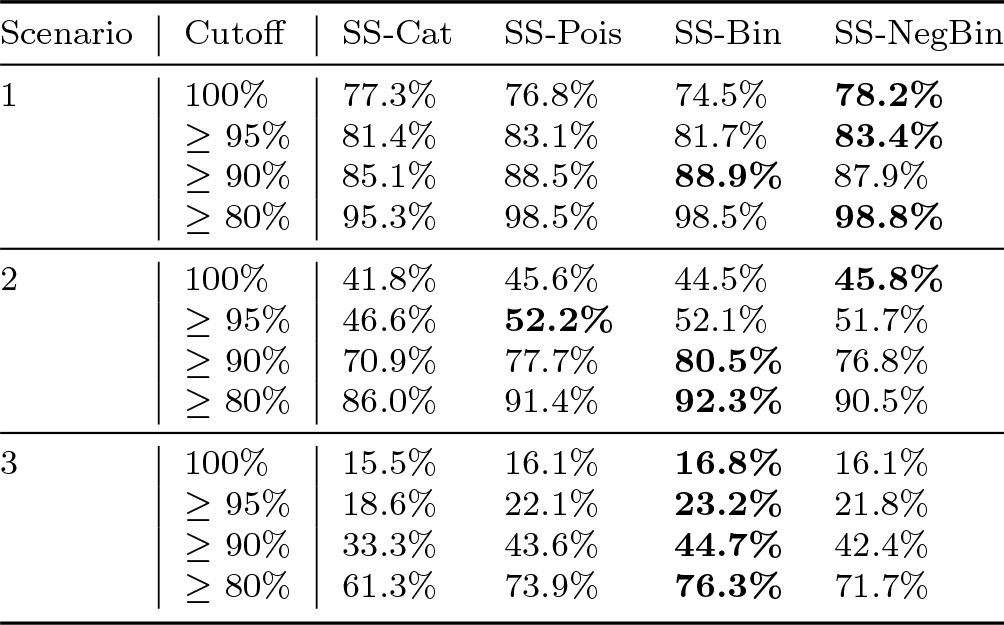
The percentage of true strains obtaining a recovery score higher than various cutoff values. True and inferred dictionaries have been simplified by merging strains that share identical SNP type at ≥ 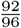 of total sites with others.

We found a positive association between the true or inferred strains’ relative strain abundance within the population and their similarity to the counterpart in the inferred or true dictionary (Figure 2). This suggests that SS constructs a haplotype dictionary that captures the frequently appearing strains in the local population. Nonetheless, the poor recovery of less abundant strains is inevitable, because there is less information to be learned.

**Fig. 2.**
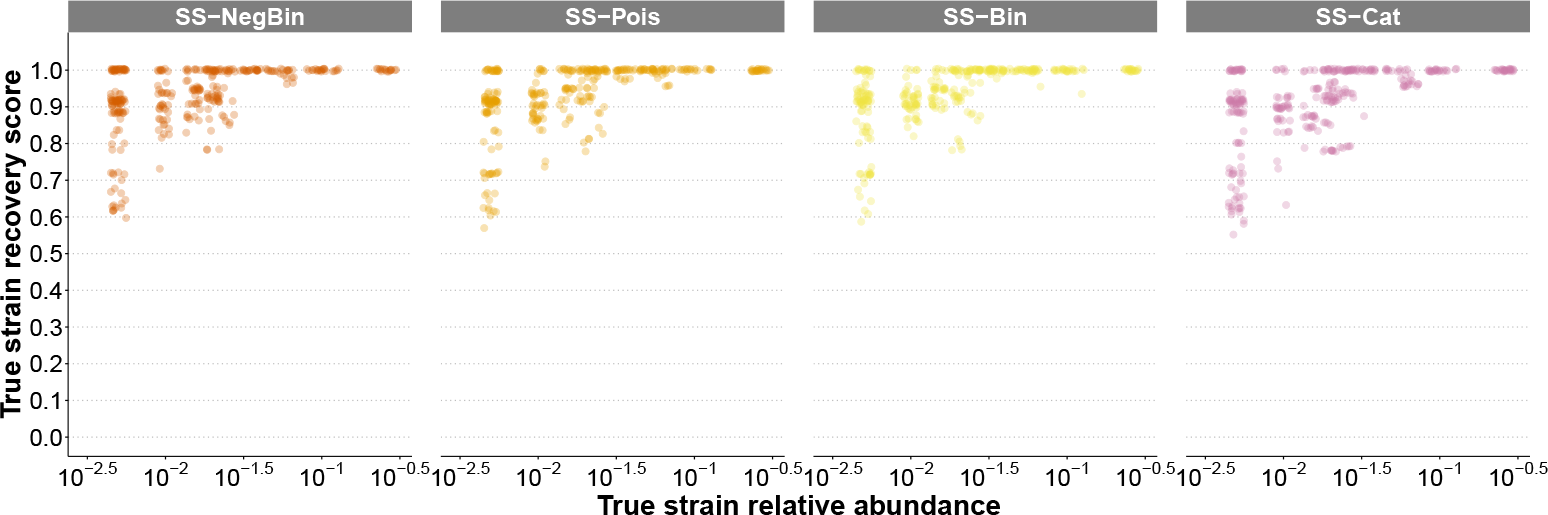
The recovery scores of true haplotypes and their relative abundance in the population. For visual clarity, a random subset of 5% points in each case is shown.

Similarly, when we inspected the similarity score of inferred strains with the definition of the largest similarity between this strain and any true strain: sim score_*l*_ = max_*k*_ sim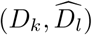, we observed a positive correlation between learned haplotype quality and their learned haplotype frequency: among the learned strains, strains with higher estimated frequencies are also more accurate (Fig. S3). Together, these suggest that SS-NegBin learns both the haplotype dictionary and strain frequencies with high fidelity.

### 2.4 SNP-Slice is robust to heterogeneous SNP sampling efforts

On empirical datasets, average read counts from genomic sequencing might differ substantially between isolate samples, especially from host tissue or blood samples, because the amount of parasite DNA and quality contained in these samples vary significantly. Signal intensities from microarray may also vary significantly, in which certain samples have weak signals. Here we tested whether SNP-Slice is robust to situations when certain samples have very low total read count information or weak SNP signal intensity. We simulated observations where a certain percentage of the hosts have only 1/10 of the total reads compared to the rest of the hosts, and tested whether our recommended method of SS-NegBin can still maintain its previously reported high accuracy. We compare the bias, RMSE, and RE of MOI, heterozygosity, and allele frequency for SS-NegBin and RM-Prop in Table 5. Since RM-Prop only uses SNP read ratio data, it is not sensitive to heterogeneity in sampling efforts. While heterogeneous sampling efforts can affect the performance of SS-NegBin, it still outperforms RM-Prop on all estimands. Moreover, we find that increasing the percentage of hosts with small total reads introduces a negative bias to MOI estimations and increases relative errors of all estimations. This is unsurprising since the quality of our inference results should deteriorate as genomic sequencing effort or SNP signal intensity decreases.

**Table 5.** Comparison of SS-NegBin and RM-Prop accuracy, when some hosts have small total read counts.

|  | 10% low reads |  | 25% low reads |  | 50% low reads |  |
| --- | --- | --- | --- | --- | --- | --- |
|  | SS-NegBin | RM-Prop | SS-NegBin | RM-Prop | SS-NegBin | RM-Prop |
| MOI |  |  |  |  |  |  |
| Bias | **0.07** | -0.65 | **-0.09** | -0.66 | **-0.25** | -0.66 |
| RMSE | **1.11** | 1.74 | **1.21** | 1.73 | **1.23** | 1.76 |
| RE | **22.4%** | 54.6% | **26.5%** | 54.0% | **27.4%** | 56.3% |
| AF |  |  |  |  |  |  |
| Bias | **-0.016** | -0.047 | **-0.023** | -0.048 | **-0.024** | -0.048 |
| RMSE | **0.030** | 0.092 | **0.041** | 0.092 | **0.043** | 0.092 |
| RE | **6.67%** | 62.9% | **12.5%** | 63.5% | **14.0%** | 63.8% |
| Heterozygosity |  |  |  |  |  |  |
| Bias | **0.002** | × | **0.002** | × | **0.002** | × |
| RMSE | **0.013** | × | **0.015** | × | **0.018** | × |
| RE | **1.608%** | × | **2.157%** | × | **3.167%** | × |

### 2.5 Empirical data evaluation on malaria in Uganda

We applied SNP-Slice to the cross-sectional survey data in Uganda to illustrate the performance of SNP-Slice on malaria field data. This survey dataset, previously investigated in [22, 33] with the RM-Cat method, contains the Sequenom assay of 105 SNPs from dried blood samples collected from random households of three sub-counties in Uganda. Due to prevalent missing data, the final included loci (*L*) and individuals (*N*) per sub-county are: Nagongera, 63L/462N; Walukuba, 49L/48N; Kihihi, 52L/74N. We re-analyzed the categorical data indicating heterozygosity status (from [22]) using the RM-Cat algorithm and compared its performance with SS-Cat. Using the same setup described in [22], we obtained the same empirical distribution of inferred MOI from RM-Cat as in Fig. 2 of [22]. RM-Cat suggests that Nagongera and Walukuba have a long and rugged tail of high MOIs with a median MOI of 5, whereas Kihihi has the highest proportion of hosts with MOI = 1, and a median MOI of 1, despite a small proportion of MOI *>* 6 (Fig. 3). SS-Cat also suggested larger proportions of mixed infections from Nagongera and Walukuba (median MOI: 3) than Kihihi, where single-infection dominates (Fig. 3). However, the distributions of MOI inferred from SS-Cat are much more concentrated, right-skewed, smooth, and do not have long and uneven tails. The maximum MOI inferred for Nagongera, Walukuba, and Kihihi are 11, 7, and 6 from SS-Cat, respectively, compared with 21, 16, and 9 from RM-Cat. The dramatic inference differences between the two methods may be attributed to the highly variable estimates from RM-Cat when the true MOI is larger than 3 (see Fig. 1). It is worth noting that RM-Cat is very sensitive to the user-specified value of maxMOI (upper bound of MOI), while inferences from SS-Cat do not require knowing the maximum MOI value a priori and are not sensitive to the value of concentration parameter *α* (Table S2). When we reduced the maxMOI value from 25 (used in [22]) to 15, the distributions of RM-inferred MOIs shifted towards the left (Figure S5). The long tail and rugged shape of the distribution, however, did not change. Given that the data includes all age classes, immunity gradually increases with age, and transmission is uniform in these localities, MOIs are expected to follow a distribution similar to either Poisson or negative binomial distributions [34]. We are able to observe this phenomenon using SS-Cat analysis but not with RM-Cat results. Therefore, SNP-Slice yields a more realistic estimation for the distribution of individual MOIs in all three regions.

**Fig. 3.**
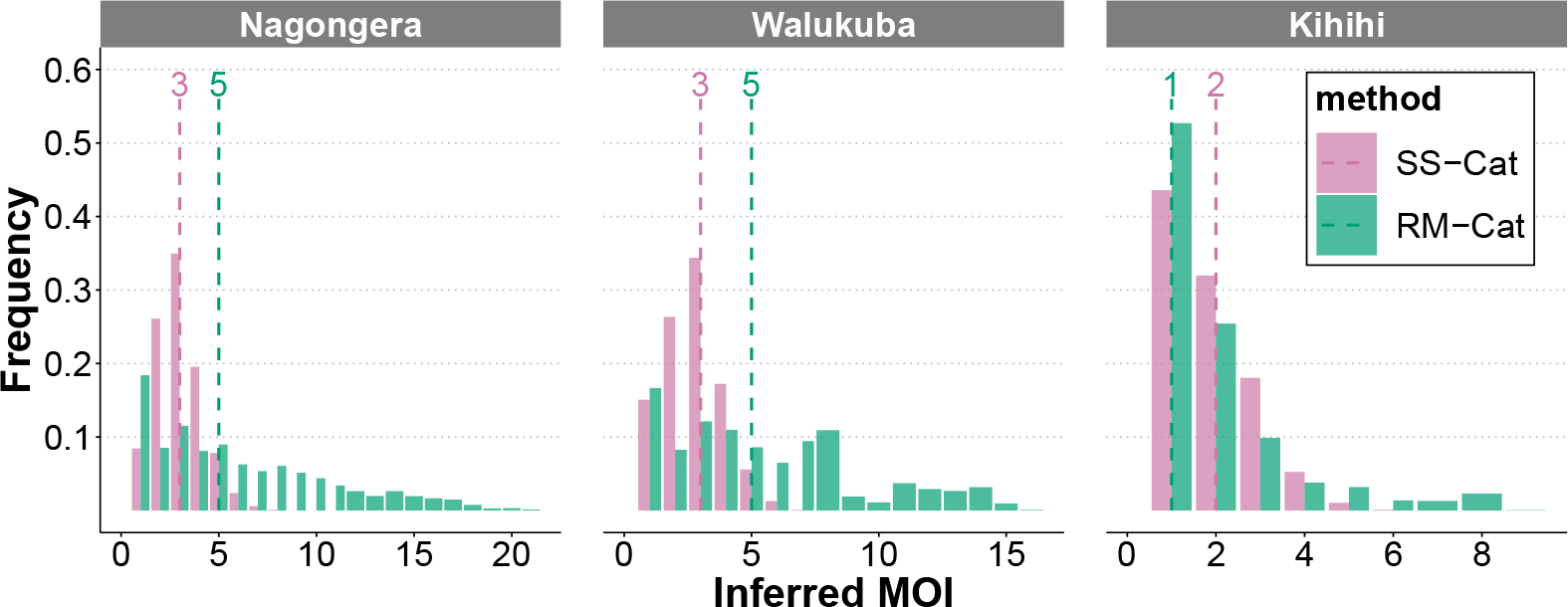
Comparison of individual MOI in three regions of Uganda estimated by RM and SS. The dotted vertical line indicates the median of the estimated MOI.

### 2.6 Empirical data evaluation on HIV

We applied SNP-Slice to longitudinal HIV-1 infection data, obtained from [35]. Unlike malaria, strain diversity in HIV does not arise from multiple infections or co-transmitted multiple strains. Instead, HIV has an extremely high mutation rate (0.34 mutations per replication cycle [36]) and forms new strains as viral load increases during the entire infection period. The constant appearance of new strains initiates an arms race with the immune system and significantly prolongs the duration of infection. Although SNP-Slice initially designed to analyze cross-sectional data, it is still applicable for such type of disease to infer the within-host strain compositions at different time points.

Here we inferred the strain dynamics within individual patients using SS-NegBin. SS-NegBin learned not only the changing pattern of MOI, the strain haplotype dictionary, but also when each strain appeared and disappeared in the evolutionary history of each patient. The number of strain haplotypes is usually low in the early stage of infection, but increases as the disease progresses (Figure 4 Top). As indicated by the lengths of horizontal line segments, some ancestral haplotypes persist throughout the tracked infection years, while most newly evolved haplotypes persist for shorter periods. This agrees with the observation that reversion to ancestral HIV-1 is common because they usually represent the optimal form for the virus [35]. Also, changes in inferred heterozygosity follow a similar trend to that of viral load, indicating that our inference result is reasonable as mutants increase when the viral load increases.

**Fig. 4.**
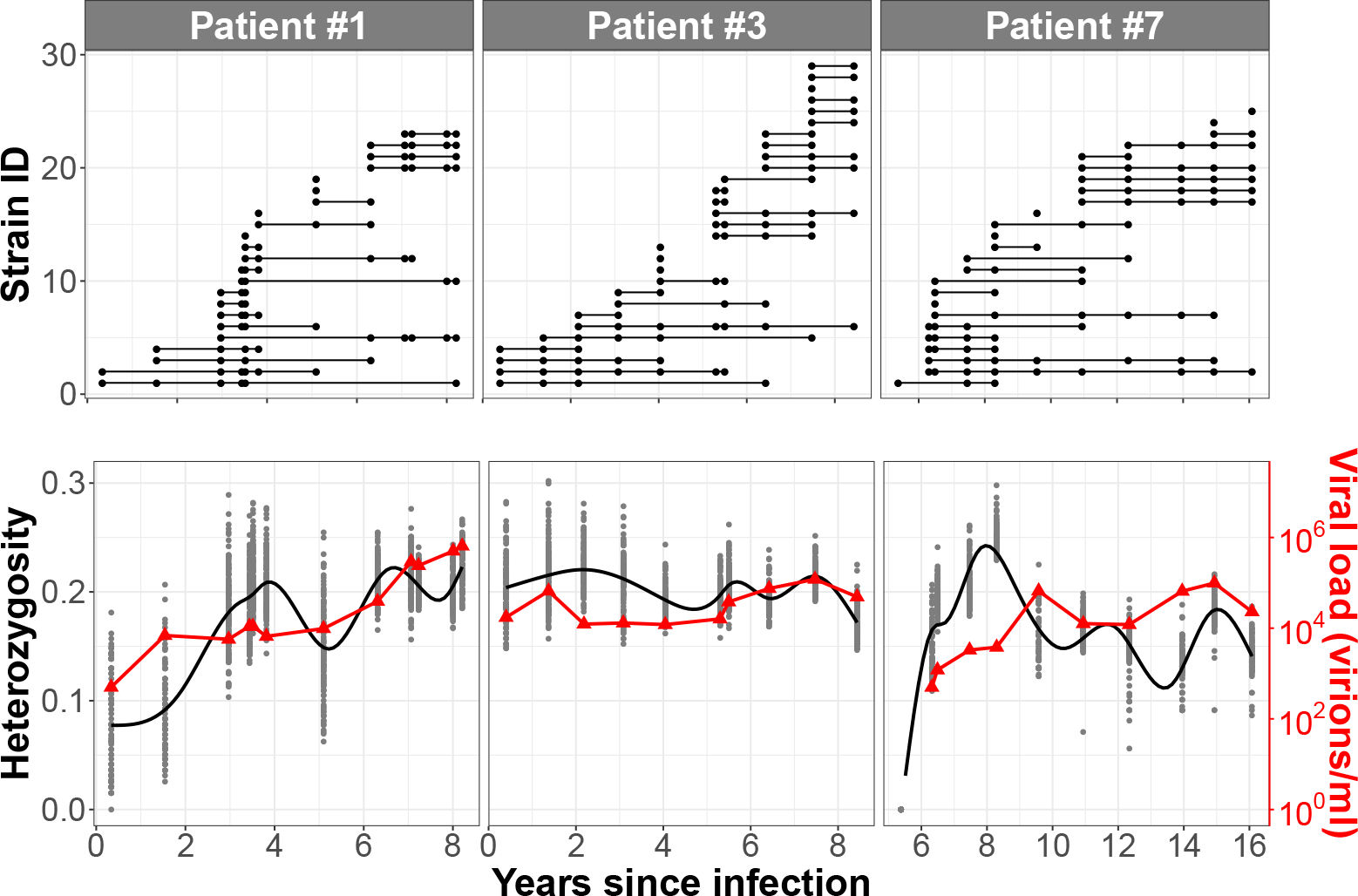
Top: appearance and disappearance of latent strains for each HIV patient. Results are based on a selected run of the SNP-NegBin method, among 128 independent repetitions, with the lowest RMSE of MOI from the average inferred MOI across all repetitions. Bottom: level of heterozygosity (left axis, gray dots and black curve) and viral load (red line, right axis). The black curve is a smooth curve connecting the averages of 128 independent repetitions of the SNP-NegBin algorithm.

## 3 Discussion

Our method is a parsimonious, effective, and reliable tool to analyze SNP read count data in the presence of mixed infections, because it 1) requires only one model parameter, 2) consistently achieves high estimation accuracy on important epidemiological quantities of interest across infection scenarios, 3) does not require reference panels. SNP-Slice has several key advantages over competing methods [20–22, 24]. First of all, in terms of unveiling novel biological insights, SNP-Slice is able to accurately reconstruct both the SNP haplotype dictionary and strain frequencies in a local population with hundreds of sites and individuals (cf. [20]). Our experiments use *L* = 96 sites and *N* = 200 −400 individuals. Despite the large size, since SNP-Slice can adapt its complexity to the data, the algorithm only needs to track a haplotype dictionary with size *<* 170, contrasting with the procedure in [20], which would operate on the full dictionary with size 2^96^. In addition to scalability, SNP-Slice is robust to unbalanced strain frequencies and variability in sampling efforts. As a result, our tool enables the estimation of heterozygosity and *population-level* strain diversity, which has not been achieved by previous methods. In particular, our haplotype dictionary and the associated strain frequency estimations can serve as a launching pad for future molecular surveillance in the same population or an adjacent region. More importantly, in terms of computational reliability, SNP-Slice *requires only one modeling parameter α*, which controls individual-MOI as well as the size of the haplotype dictionary, and the algorithm is *robust* to this user-specified value (Section S1.4). In contrast, DEploid [19]) imposes an upper bound on the number of within-host haplotypes; RM requires several input parameters and demonstrates sensitivity to `maxMOI` (Section S1.5).

Here we make some recommendations for using SNP-Slice. When estimating regions with prevalent mixed infections, e.g. average MOI *>* 2, one should use read counts data instead of heterozygosity status data if possible, as the former contains more detailed mixed infection information for resolving higher MOIs. Our experiments have confirmed that count-based methods are more accurate than SS-Cat, with the only exception of estimating MOI in the low transmission intensity setting (average MOI at 1.27). Our analysis suggests that SS-NegBin is preferred for studying field samples, where the observation noise can be high. Even in the most challenging scenario, *>* 75% of the MOI estimations from SS-NegBin are within ±1 of the true value and relative error *<* 29%. SS-Pois performs as well as SS-NegBin in our simulated data when transmission intensity is high (Scenario 3), but might also be attributed to that we use Poisson distribution to simulate read counts (Section S2.3). Since SS-NegBin is designed to be more robust to violations of model assumptions (Section 4.2), it is still the recommended method within the SNP-Slice family.

Our re-analysis of the Uganda malaria SNP array data differs significantly from the original analysis in [22], but roughly agrees with another study that sequenced the major antigen gene *var* DBL*α* types [37]. Unique DBL*α* types sampled from Nagongera, Walukuba, and Kihihi were, on average, 118, 98, and 68 in [37] respectively. Each parasite genome usually carries around 50-60 unique *var* genes. Because more than 75% of types only appeared once in these populations, we can safely assume each strain carries around 40 unique types, translating to a mean MOI of 3.0, 2.5, and 1.7 in Nagongera, Walukuba, and Kihihi. The maximum MOI in Nagongera and Walukuba does not exceed 5.3 from the *var* type data, or 6-7 from SS-Cat, while using RM-Cat would lead to the conclusion that many individuals have MOI larger than 10, which is unrealistic given the transmission intensity in the region.

Although SNP-Slice is designed to resolve mixed infections in a local population, it can also be applied directly to a longitudinal sample of strain evolution within a host [35, 38]. When applied to the time series of HIV-1 within patients, SS recovered haplotype dynamics in HIV and achieved reasonable performance. Zanini et al. [35] showed that the phylogenetic tree of V3 region built from patient 1 displays one low haplotype diversity period between two episodes of bursts of new haplotypes, while Patient 3 has a more steady rate of strain replacement. We recovered a similar pattern in Figure 4 regarding the dynamics of strains.

We discuss some limitations of SNP-Slice in its current form and point out several ways to extend this framework. First, SS characterizes representative hosts carrying common haplotypes in the population better than atypical hosts with rare haplotypes, for example, those hosts investigated in Section S1.3 and rare haplotypes in Figure 2. This is unsurprising because SS pools information across hosts to improve haplotype inferences, hence favoring common haplotypes and typical hosts. Following this logic, SS is intended to study mixed infections in a local population. We do not recommend using SS on artificially pooled samples from different localities unless there is enough overlap in the haplotype dictionaries across these populations. Second, because the SNP-Slice framework reconstructs the haplotype dictionary from scratch, it requires that the number of hosts in the dataset be large enough to contain the most abundant strains in the local population. While SS is not intended to analyze small datasets or single host information, if necessary, the framework can be extended to incorporate existing haplotype reference panels so that one can apply the method separately to each host. Finally, we emphasize that SNP-Slice is a modeling framework and is not limited to the four methods (Bin, Pois, NegBin, and Cat) studied in this article. Users can choose their own observational models suitable for their experimental protocols. For example, if rounds of PCR are used to generate amplicons before sequencing, then a user can incorporate more overdispersion beyond the current models presented in Section 4.2.

In summary, we developed a general methodology for resolving mixed infections on hosts infected by a population of parasites. SNP-Slice can be widely applied to any pathogen SNP panels as long as they contain biallelic markers and have information on read count, SNP intensity signal, or heterozygosity status. The resolved haplotype information recovered from polygenomic data by SNP-Slice enables many standard analyzing tools for understanding genomic relatedness, diversity patterns, and evolutionary trends. Since SNP-Slice can accurately estimate host-level heterozygosity and construct population-level haplotype dictionaries, it is expected that SNP-Slice will become a standard tool for disease surveillance in molecular epidemiology.

## 4 Methods

### 4.1 Problem formulation

Our data consists of SNP read counts (or heterozygosity status) for *N* hosts at *L* sites. For each host *i* and each site *j*, we observe 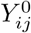 reads from major alleles and 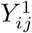 reads from minor alleles, summing to 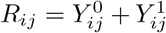 total reads. When read counts data is not available, one observes heterozygosity status at each site (0: only major alleles, 1: only minor alleles, 0.5: heterozygous).

SNP-Slice aims to resolve mixed infections by estimating two matrices: the feature dictionary *D* and a feature assignment matrix *A*. The assignment matrix *A* ∈{0, 1}^*N*×*K*^ describes which strains are present in each host: *A*_*ik*_ = 1 indicates the *i*th host is infected with the *k*th strain. The feature dictionary *D* ∈ {0, 1}^*K*×*L*^ represents the underlying strains at the population level. Each row of *D* represents the haplotype of one strain: *D*_*kj*_ = 0 indicates the *k*th strain carries a major allele at site *j* and *D*_*kj*_ = 1 indicates the site has a minor allele. For the haplotype dictionary, we choose independent Bernoulli priors *D*_*kj*_ ∼ Bernoulli(*ρ*). In our experiments, we have set 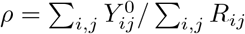 for SNP counts data and *ρ* = 0.5 for categorical data.

The size of the haplotype dictionary, *K*, is usually unknown and determines the complexity of the model. All length *L* binary sequences are candidates for the haplotype dictionary, so there are *K* = 2^*L*^ ≈ ∞ potential haplotypes. SNP-Slice takes a Bayesian nonparametric approach [39, 40], and thus allows the complexity of the model to grow as more data are observed.

### 4.2 Three novel noise models tailored to SNP counts

We describe observational models linking the latent variables (*A, D*) describing mixed infection structure and to the mixed infection observations *Y*. SNP-Slice uses the categorical model proposed in [22], when we only observe heterozygosity status.

Here we propose three novel noise models tailored to SNP read counts data. Given *A* and *D*, a host *i* has 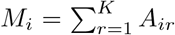 strains, and here *M*_*i*_ is the MOI of host *i*. At each site *j*, this host should have 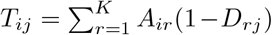 major alleles. Treating total read counts *R*_*ij*_ as fixed, let’s assume 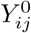’s are mutually conditionally independent given *A, D*, across hosts and sites. To begin with, we introduce a Binomial model, where

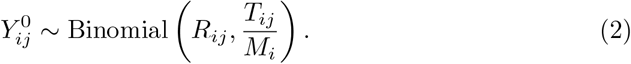

Under the Binomial model, for each *i, j*, all the *M*_*i*_ strains infecting host *i* has the same chance of appearance among the *R*_*ij*_ total SNP reads. Due to Binomial model’s strong adherence to the ‘balanced weights’ assumption and because the actual frequency of each strain within the host is usually uneven and varies temporally, we also propose two more flexible models: the Poisson model with

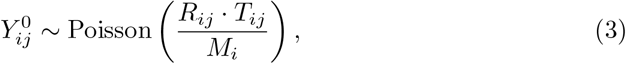

and the Negative Binomial model

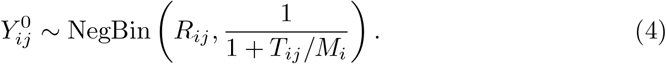

The Poisson and NegBin models progressively introduce higher variance to *Y* ^0^ while holding *R, A, D* fixed (see Table S3), and therefore should be more robust to model miss-specifications such as balanced strain weights during observation. This is consistent with our empirical evaluations in Section S1.2. In Section 2.4, we also confirmed that the NegBin model is robust against heterogeneous sampling efforts across hosts, which is frequent in field samples.

### 4.3 Bayesian nonparametric inference with the IBP prior

As we uncover latent strain structures behind the observed SNP data, a key challenge lies in determining an appropriate level of model complexity, as reflected in dictionary size *K*. SNP-Slice takes a Bayesian nonparametric [28, 40] approach to model selection. Instead of specifying a fixed number of strains in the population or an upper bound in it, SNP-Slice can learn *K* from the empirical data, allowing the model complexity to grow as we observe more data. Additionally, since in mixed infections, each host can be infected with any combination of strains, our model has to account for such a combinatorics structure.

The considerations above lead us to choose the India Buffet Process prior for *A*. The IBP is an infinite dimensional probability distribution over sparse binary matrices with a finite number of rows (representing hosts) and an unbounded number of columns (representing features/strains). SNP-Slice only uses one parameter *α*, which controls the number of unique strains in a finite sample as well as the sparsity of the matrix *A*. Like many prior distributions used in Bayesian nonparametrics modeling [28, 40], the IBP [26, 28, 31] is inspired by a culinary metaphor: Consider a restaurant that serves infinitely many dishes, each of them representing one feature/strain, where customers (hosts) eat existing dishes based on their popularity and try Pois(*α*) numbers of new dishes. This representation is conducive to understanding the clustering structure in feature assignments, where popular features tend to become more popular as the number of hosts increases. More importantly, the number of dishes tried by each customer, analogous to MOIs in our data, should follow a Pois(*α*) distribution, consistent with prior knowledge about malaria MOIs [34]. We believe SS-Cat’s superior performance over RM-Cat in Fig. 3 can be attributed to this property of the IBP prior. However, here we present a stick-breaking representation of the IBP [25] that facilitates our MCMC computations; See Fig. 5 for an illustration. Let *µ*_(1)_ *> µ*_(2)_ *>* … *> µ*_(*K*)_ be a decreasing ordering of (*µ*_1_, *µ*_2_, …, *µ*_*K*_) where each 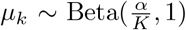 independently. Let *A* be a binary matrix of size *N* × *K*, where *A*_*ik*_ ∼ Bernoulli(*µ*_(*k*)_).

**Fig. 5.**
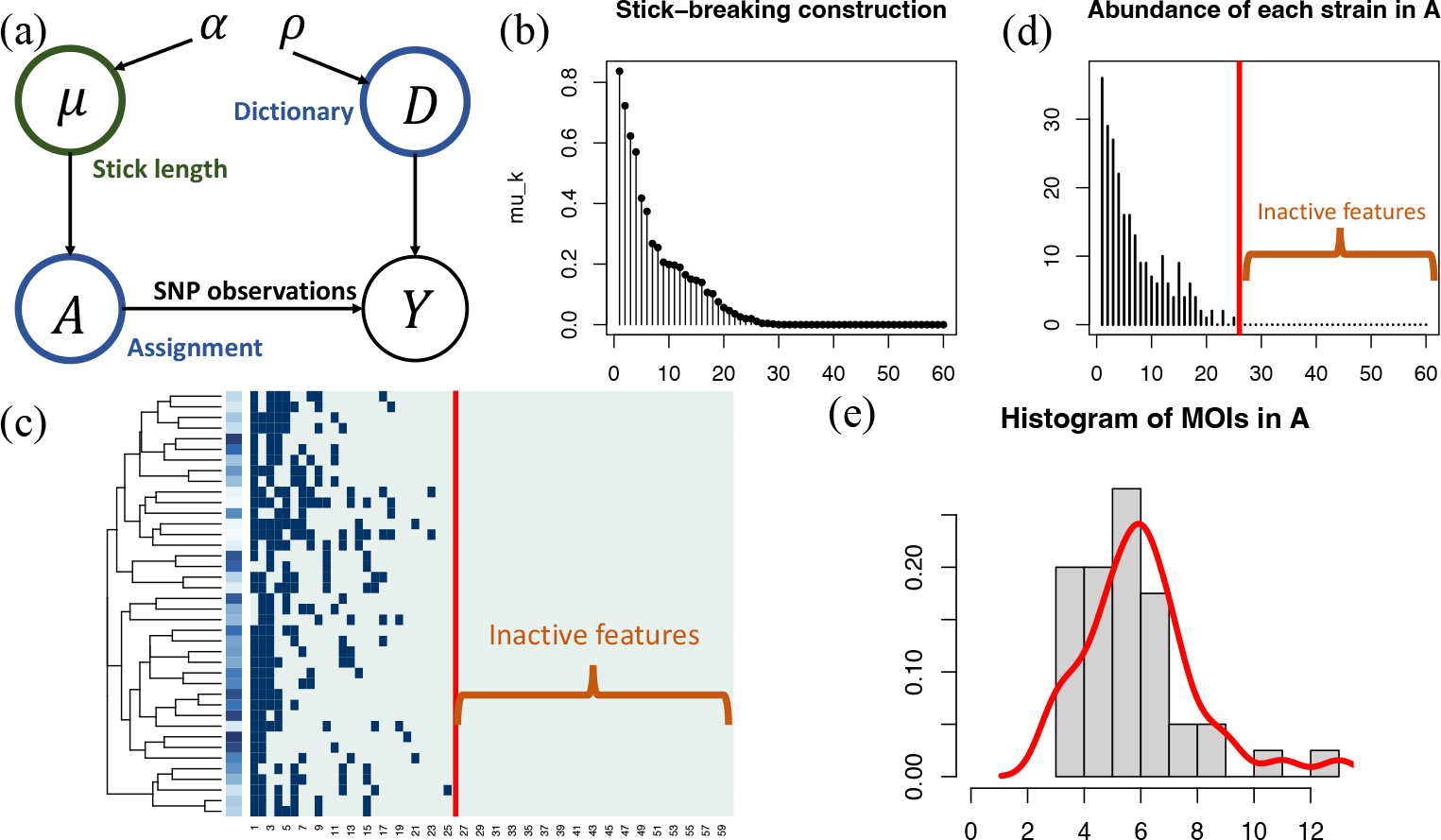
Illustration of the stick-breaking construction of the Indian Buffet Process. (a) A graphical model representation of the SNP-Slice model. *α* and *ρ* are model parameters, *µ, A, D* are latent variables, and *Y* is observed. (b) Stick-length of the first 60 strains (ordered by *µ*_*k*_ values, with *α* = 5. (c) In a random sample of *A* from IBP, each row represents a host and each column corresponds to a strain. The strains have been clustered based on their co-appearance structure, and strains from #26 onwards are inactive strains. (d) The abundance of the first 60 strains, following a similar shape with the stick lengths in (b), but only have *K*^+^ = 25 active features. (e) Histogram of host MOIs and fitted density curve. In theory, they should follow a Pois(5) distribution.

So far *α* is a concentration parameter and *K* is a truncation level that corresponds to the unknown dictionary size. In the limit of *K →* ∞, we have *A* ∼IBP(*α*). The stick-breaking metaphor begins with a stick of unit length. One recursively breaks a length *µ*_(*k*)_ stick into two smaller pieces at the *ν*_*k*_ ∼ Beta(*α*, 1) relative point, and the next headpiece shall have length *µ*_(*k*+1)_ = *µ*_(*k*)_*ν*_*k*_. The length of *µ*_(*k*)_ shall correspond with the frequency of the *k*th most popular feature in *A*. As the stick-breaking process continues, the remaining stick will be so small that all subsequent features are inactive. While we can continue the process until *K →* ∞, there should only be a finite number of active features, denoted as *K*^+^. For example, the IBP sample in Fig. 5 has only *K*^+^ = 25 active features.

### 4.4 The SNP-Slice algorithm

We develop an SNP-Slice algorithm to approximate the posterior distribution of *A, D*. The algorithm is based on slice sampling [25, 41], which is a class of Markov Chain Monte Carlo algorithm [42, 43] that can handle complex distributions, including the posterior distributions induced by the three non-conjugate observation models tailored to SNP read counts data. SNP-Slice utilizes the stick-breaking construction for the IBP, which includes stick lengths *µ* as an auxiliary variable. The sampling problem is challenging, because all three latent variables are infinite-dimensional: *µ* has infinitely many entries, *A* has infinitely many columns, and *D* has infinitely many rows.

Following [25], we facilitate computations by introducing an auxiliary slice variable

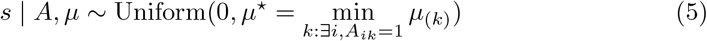

where 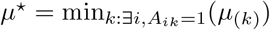 is the shortest stick length of all active features. Importantly, this construction guarantees that *s* is less than the stick length corresponding to any active feature, so those features associated with lengths of *µ*_(*l*)_ *< s* must be inactive. As a result, knowing *A*_*il*_ = 0 for all *µ*_(*l*)_ *< s*, we can fix the parts of (*A, D*) corresponding to inactive features *µ*_(*l*)_ *< s*. After adding *s*, the state space becomes {*µ, A, D, s*}. SNP-Slice proceeds by updating these variables iteratively while keeping the joint posterior distribution invariant. Our implementation maintains a finite number of sticks to circumvent the curve of infinite dimensionality and it increases the dimensions of *µ, A, D* when needed.

The slice variable *s* determines this finite truncation level, which adapts to the complexity of the data. When *s ≤µ*_(*k*)_ for all features, we will break more sticks to create an inactive feature, sampling according to a distribution with (unnormalized) probability density function

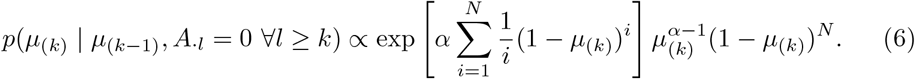

We only need one feature with *µ*_(*k*)_ ≤*s*, since the construction forces all subsequent features to also be inactive.

Potentially active features (*µ*_(*k*)_ *≥ s*) each have infected 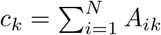 hosts. The conditional distribution of *µ*_(*k*)_ follows the Beta(*c*_*k*_, *N* + 1 *− c*_*k*_) distribution restricted to the interval (*µ*_(*k*+1)_, *µ*_(*k−*1)_). Since features with *µ*_(*k*)_ *< s* are definitely inactive, we can update their stick lengths using Eq.(6). Since entries in *A* and *D* are binary, their conditional distributions are Bernoulli distributions. We do not need to update parts of *A* and *D* corresponding to inactive features. Details are discussed in Section S2.2.

### 4.5 Epidemiological Quantities Derived from SNP-Slice Results

Many genetic and epidemiological quantities can be derived from SNP-Slice results to benefit molecular biology research and public health decisions. In Section 2, we used MOI, AF, and host-level heterozygosity to assess the accuracy of our methods.

At the population level, one can calculate minor allele frequencies per locus with 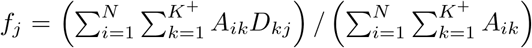. The corresponding major allele frequency is *F*_*j*_ = 1 *− f*_*j*_. Heterozygosity in the local parasite population *H*_*S*_ can be calculated with 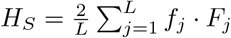.

For each host, one can calculate MOI 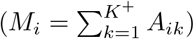, heterozygosity (*H*_*i*_), and *F*_ws_. Within host heterozygosity [44], *H*_*i*_ ∈ [0, 0.5], reflects the average level of allelic diversity over all the sites. It can be computed from SNP-Slice outputs with

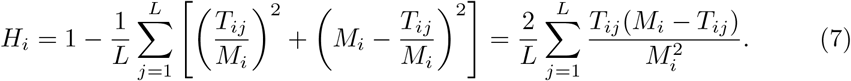

A single-infection host has *H*_*i*_ = 0. Multi-strain infections have *H*_*i*_ *>* 0, and heterozygosity is the highest (=0.5) when a host has two strains, each with frequency 50%, that are exactly opposite of each other. *F*_WS_ [45] is another metric that reflects the complexity of infection in each host. It can be calculated with *F*_*WS*_ = 1 *− H*_*i*_*/H*_*S*_.

## Acknowledgments

We thank Dr. Frédéric Labbé for assistance in utilizing the malaria-SNP simulator to generate simulated SNP read counts data. Q. He and N. Ju acknowledges support from Purdue University’s Showalter Trust Research Award. Also, this project was funded, in part, with support from the Indiana Clinical and Translational Sciences Institute funded, in part by Grant Number UL1TR002529 from the National Institutes of Health, National Center for Advancing Translational Sciences, Clinical and Translational Sciences Award. The content is solely the responsibility of the authors and does not necessarily represent the official views of the National Institutes of Health.

## Author Contributions

N. Ju designed, coded, and tested the SNP-Slicer algorithm. J. Liu analyzed empirical and simulated data. N. Ju and Q. He jointly conceived and supervised the work. N. Ju wrote the first draft. All authors revised the manuscript.

## Competing Interests

The authors declare no competing interests.

## Supplemental Materials

### S1 Supplementary Results

#### S1.1 Additional results on the accuracy of SNP-Slice

In Section 2.1, we compared the overall bias, RMSE, and RE of AF estimations between SS and RM numerically in Table 2. Here we inspect whether the bias and RE change according to the true allele frequency. As shown in Figure S1 of Scenario 2, the average error of SS-Bin, SS-Pois, and SS-NegBin stay around −0.01 along the entire range of AF, which is at least 1 magnitude smaller than the true AFs if rare SNPs are filtered before the analysis. This is also reflected by the small relative errors in Table 2. In contrast, RE increases with higher true AF in RM-Prop.

**Fig. S1.**
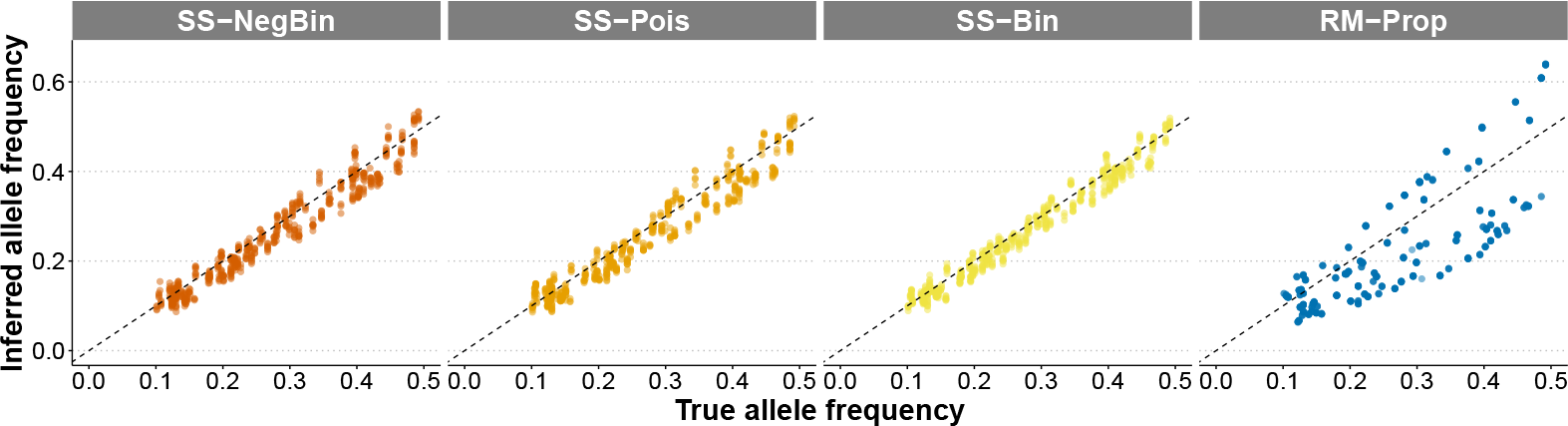
True allele frequency and their estimations from SS-NegBin, SS-Pois, SS-Bin, and SS-Cat. A randomly selected subset of 5% of the points are displayed for visual clarity.

When comparing the heterozygosity estimation between read counts vs. categorical methods, we observe that the error of heterozygosity estimations in read-count based SS is uniform along the true heterozygosity (Figure S2), while SS-Cat has a strong positive bias when the true heterozygosity is above 0.2. This aligns with the fact that SS-Cat tends to underestimate strain diversity in hosts with high MOIs.

**Fig. S2.**
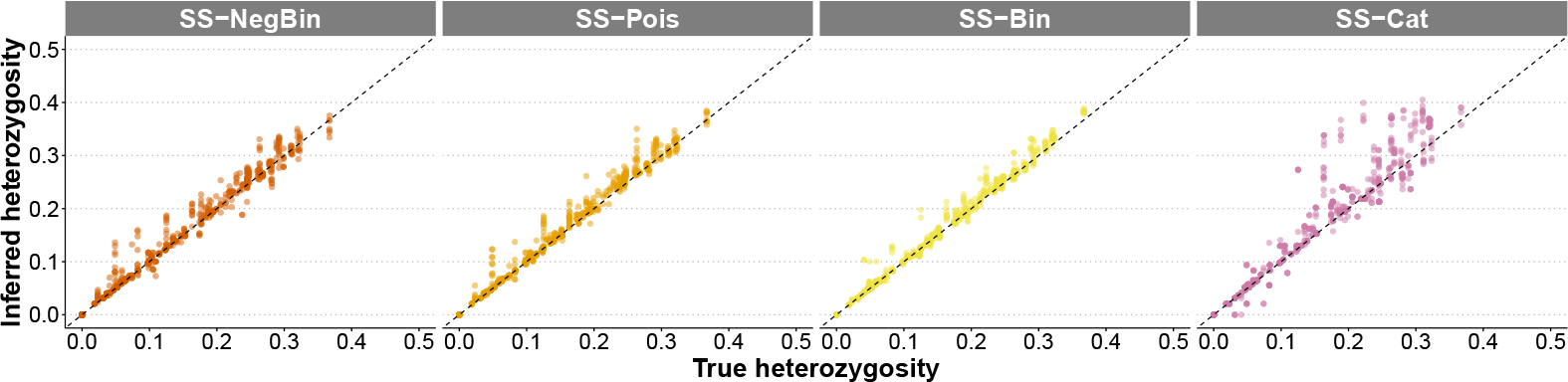
True heterozygosity and their estimations from SS-NegBin, SS-Pois, SS-Bin, and SS-Cat. A randomly selected subset of 5% of the points are displayed for visual clarity.

Figure S3 displays the positive correlation between the quality of inferred haplotypes (measured with sim score_*l*_ = max_*k*_ sim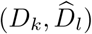 and their relative abundance in 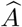.

**Fig. S3.**
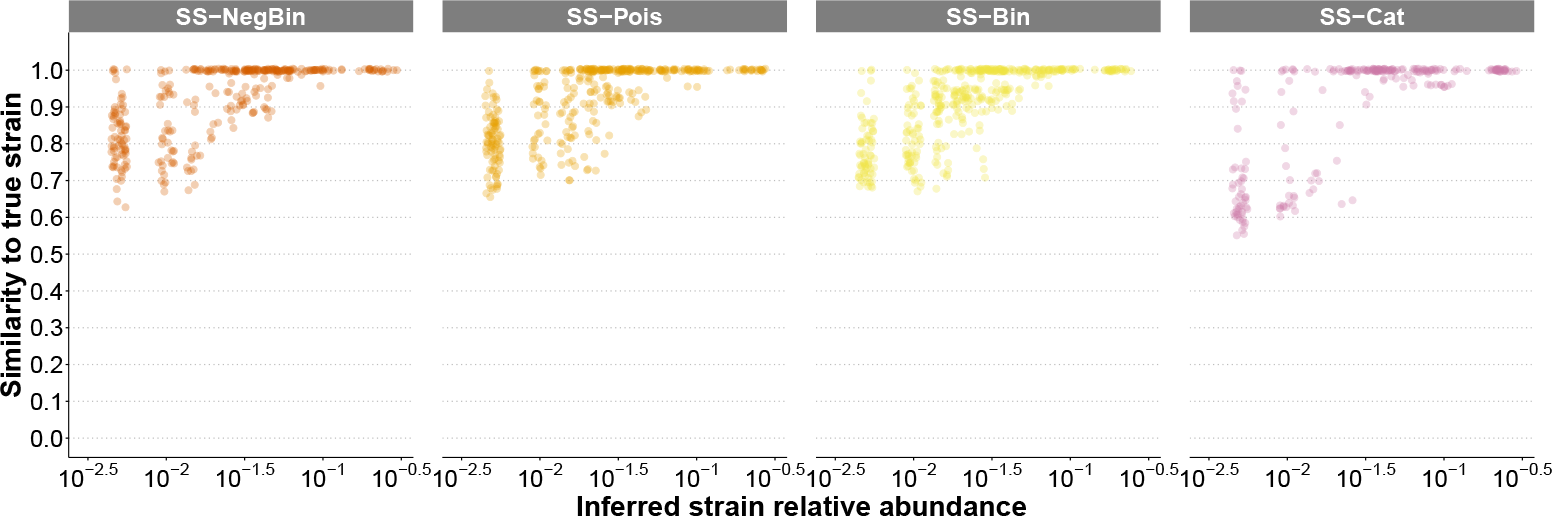
The similarity of inferred haplotypes and their relative abundance in the population.

#### S1.2 SNP-Slice is robust to unbalanced within-host strain frequency

When modeling SNP read count data, SS-Bin, SS-Pois, and SS-NegBin make the assumption that all strains within the same host appear with equal frequency. This modeling assumption facilities efficient computations. However, within-host competition of strains due to the difference in growth rates or drug sensitivity often leads to unbalanced composition of strain frequencies. We tested the robustness of the current model to the violation of this balancing assumption, by simulating an SNP read count dataset where strains within the same host have unbalanced strain frequencies. SS-NegBin still outperformed RM-Prop when estimating MOI, AF, and heterozygosity. Also, comparing Table S1 with the bias, RMSE and RE reported for Scenario 2 in Tables 1,2, and 3, the performance of SS-NegBin does not deteriorate when the balanced-frequency assumption is violated. This is because SS-NegBin is designed to be more robust to violations of model assumptions, as we will explain in Section 4.2.

#### S1.3 Potential cause of bias in MOI due to minor allele sites

In Section 2.1 and Figure 1, we see that both SNP-Slice and RM makes larger mistakes on large MOI individuals, as indicated by the wide boxes. In this section, we investigated a potential cause of these large biases in MOI estimation by both methods. It is worth noting that although major alleles should appear more frequently than minor alleles in the population, there can be some hosts that have more minor allele sites than major allele sites. Our investigations in Figure S4 focus on these atypical hosts. In every host, we can mark each site as minor-dominated or major-dominated based on its haplotype compositions. For hosts with *<* 20% minor-dominated sites, all methods control estimation error below ±1. When the fraction of minor-dominated sites is around 40%, the bias of RM-Prop is around −3; in contrast, the average error from our SS-based method is around 0,1 and 2 for SS-NegBin, SS-Pois, and SS-Bin respectively. The increase in MOI estimation bias is manifested as the fraction of minor-dominated sites increases above 40%: SS-NegBin, SS-Pois, and RM-Prop underestimates MOI while SS-Bin overestimates MOI.

**Table S1.** Comparison of SS-NegBin and RM methods, under unbalanced within-host strain frequencies.

|  | SS-NegBin | RM-Prop |
| --- | --- | --- |
| MOI |  |  |
| Bias | **0.36** | -0.74 |
| RMSE | **1.54** | 1.82 |
| RE | **43.1%** | 59.7% |
| AF |  |  |
| Bias | **-0.010** | -0.050 |
| RMSE | **0.028** | 0.093 |
| RE | **5.89%** | 64.3% |
| Heterzygosity |  |  |
| Bias | **-0.002** | × |
| RMSE | **0.026** | × |
| RE | **6.630%** | × |

**Fig. S4.**
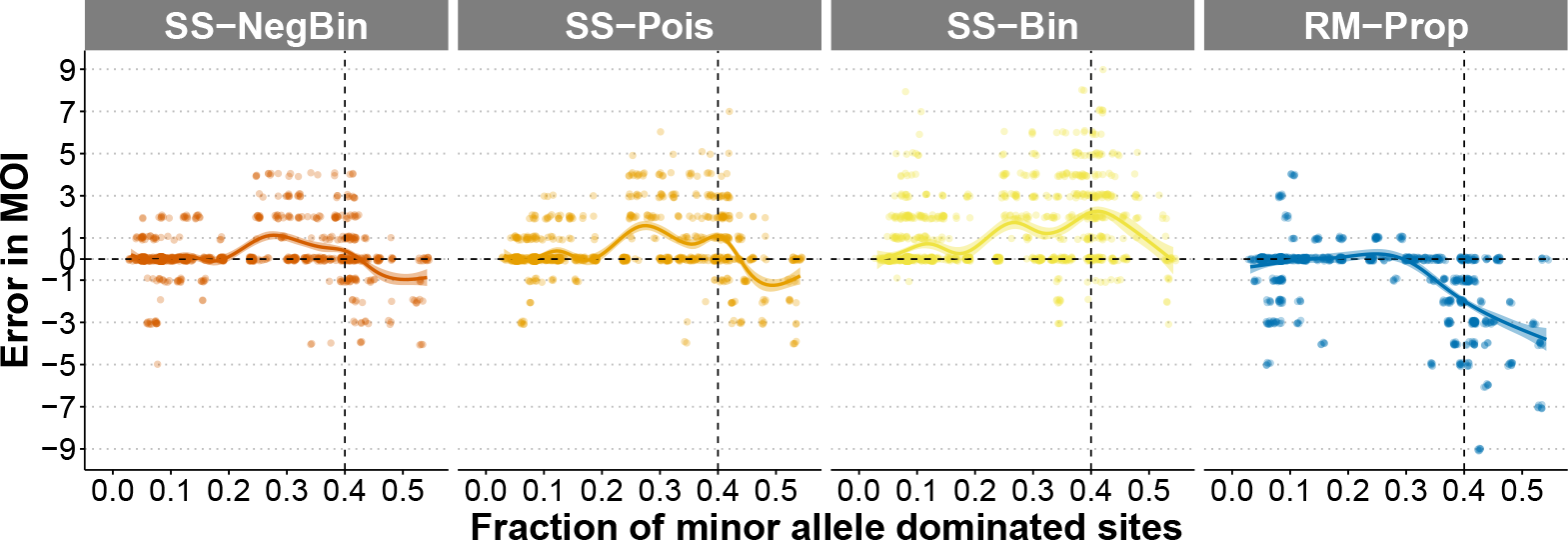
Error of MOI estimation using count-based SNP data in Scenario 2, where hosts are organized based on the fraction of sites with more minor allele reads than major allele reads. For visual clarity, a random subset of 5% points is displayed for each method.

#### S1.4 Sensitivity of SNP-Slice to *α*

In Section 2, we have shown detailed results for simulated Scenario 2, where we used *α* = 3 as the input value of the concentration parameter since the average true MOI is 3 2.58. Here we investigated the sensibility of SNP-Slice to the concentration parameter *α*. We changed *α* to 1.5 and 4, the input values for Scenario 1 and 3 respectively, and reran SS-NegBin and SS-Cat for Scenario 2. As Table S2 shows, different *α* values lead to similar output results, indicating that SNP-Slice is robust to this parameter. In other words, SNP-Slice does not require much prior knowledge of the population-level mean MOI or transmission intensity. It can be widely applied to analyze various scenarios and generate reliable inferences.

**Table S2.** Sensitivity of SNP-Slice to the value of concentration parameter *α*.

|  | SS-NegBin |  |  | SS-Cat |  |  |
| --- | --- | --- | --- | --- | --- | --- |
| | $\alpha = 1.5$ | $\alpha = 3$ | $\alpha = 4$ | $\alpha = 1.5$ | $\alpha = 3$ | $\alpha = 4$ |
| MOI |  |  |  |  |  |  |
| Bias | 0.01 | 0.10 | 0.14 | -0.57 | -0.58 | -0.58 |
| RMSE | 1.09 | 1.10 | 1.12 | 1.46 | 1.48 | 1.48 |
| RE | 21.7% | 22.1% | 22.8% | 38.5% | 39.8% | 39.4% |
| AF |  |  |  |  |  |  |
| Bias | -0.015 | -0.015 | -0.015 | -0.041 | -0.041 | -0.041 |
| RMSE | 0.030 | 0.029 | 0.029 | 0.093 | 0.092 | 0.092 |
| RE | 6.58% | 6.45% | 6.44% | 64.2% | 63.9% | 63.7% |
| Heterozygosity |  |  |  |  |  |  |
| Bias | 0.003 | 0.003 | 0.003 | 0.008 | 0.008 | 0.008 |
| RMSE | 0.014 | 0.012 | 0.011 | 0.031 | 0.031 | 0.031 |
| RE | 1.895% | 1.285% | 1.169% | 9.558% | 9.292% | 9.366% |

#### S1.5 Sensitivity of RM to maxMOI

As described in Section 2.5, the RM-Cat method generates MOI distributions with long and rugged tails. Here we change the user-specified upper bound of MOI (maxMOI) to 15 instead of 25 and rerun RM analysis (Fig. S5). Although the MOI distributions in the new setting still have long tails, we observe notable changes in the inferred MOI distribution, particularly for Nagongera and Walukuba. With maxMOI=25, RM estimated the maximum MOI to be 21 and 16 for these two populations. However, when setting maxMOI=15, the largest inferred MOI was reduced to 14 and 11 respectively. Higher transmission regions, like Noagonera and Walukuba, are more sensitive to the value of maxMOI, which impacts our estimations on high MOI individuals. Notably, SNP-Slice is not sensitive to the value of the concentration parameter *α*, which enables it to outperform The Real McCOIL in generating more realistic MOI distributions with minimal prior information.

**Fig. S5.**
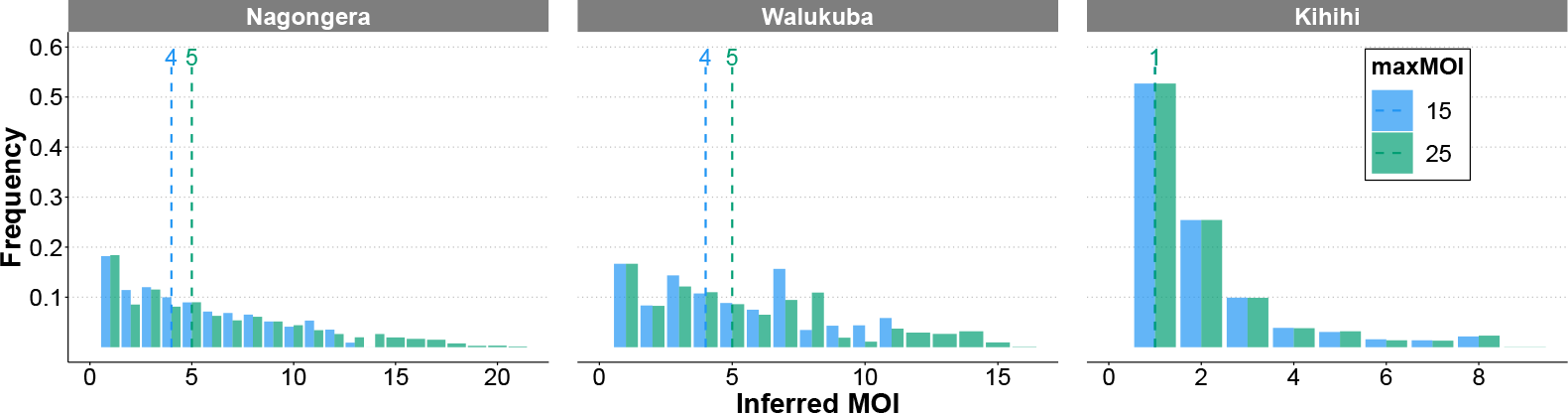
Comparison of individual MOI in three regions of Uganda estimated by RM, when maxMOI is set to 15 (blue) instead of 25 (green, used in Fig. 3). The dotted vertical lines indicate medians of the estimated MOI.

### S2 Supplementary Methods

#### S2.1 An Analytical Comparison of the Three Noise Models

The three noise models have the same expected values for *Y*_*ij*_ and the same maximum likelihood estimators (MLE) for *T*_*ij*_ when the individual MOI *M*_*i*_ is known. But in terms of variance of *Y*_*ij*_, NegBin *>* Pois *>* Bin.

**Table S3.** Comparison of three noise models.

| Model | $\mathbb{E}(Y_{ij})$ | $\text{Var}(Y_{ij})$ | MLE of $T_{ij}$ given $M_i$ |
| --- | --- | --- | --- |
| Bin | $R_{ij} \cdot T_{ij}/M_i$ | $R_{ij} \cdot (T_{ij}/M_i)(1 - T_{ij}/M_i)$ | $M_i \cdot Y_{ij}/R_{ij}$ |
| Pois | $R_{ij} \cdot T_{ij}/M_i$ | $R_{ij} \cdot T_{ij}/M_i$ | $M_i \cdot Y_{ij}/R_{ij}$ |
| NegBin | $R_{ij} \cdot T_{ij}/M_i$ | $R_{ij} \cdot (T_{ij}/M_i)(1 + T_{ij}/M_i)$ | $M_i \cdot Y_{ij}/R_{ij}$ |

#### S2.2 Additional details of the SNP-Slice algorithm

##### S2.2.1 Update *D*

The conditional posterior distribution for the dictionary is *p*(*D* | *µ, A, s, Y*) ∝ *p*(*D*)*p*(*Y* | *A, D*), which is independent of *µ* and *s*. We can iteratively update each atom *D*_*kj*_ in the dictionary with samples from the Bernoulli distribution. The two conditional probabilities are 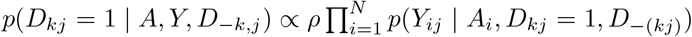 and 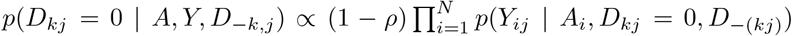. We only need to update the active features/strains, because the inactive features do not enter the likelihood *p*(*Y* | *A, D*) specified by the three observation models.

##### S2.2.2 Update *A*

Similar to updating *D*, we can iteratively update each atom *A*_*ik*_ with a sample from the Bernoulli distribution, following the conditional distribution *p*(*A* | *D, µ, s, Y*) ∝ *p*(*A* | *µ*)*p*(*s* | *A, µ*)*p*(*Y* | *A, D*). The two conditional probabilities are: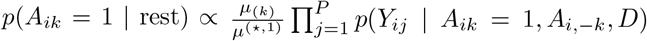 and 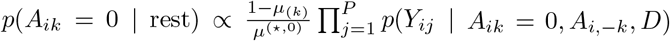. The denominators, namely *µ*^(*⋆*,1)^ and *µ*^(*⋆*,0)^, are needed when a change in allocations *A*_*ik*_ affects which feature is the last active feature.

#### S2.3 Simulated Malaria Infection Data

Our simulation experiments in Sections 2.1-2.4 uses simulated SNP data generated from an agent-based stochastic simulator of malaria transmission. The simulator tracks the evolution of parasite strain composition and host immunity (see details in [6, 32]). At the beginning of the simulation, each parasite is initialized with a randomly sampled SNP set defined by the minor allele frequency (MAF) per site.

In Section 2, we tested three different levels of transmission intensity that encompass endemic transmissions in Africa to seasonal transmissions in Asia or South America. The Scenarios 1-3 represent simulations with transmission rates from low to high. In the simulation process, we used the same parameter values from [32], assuming identical initial SNP allele frequencies and no linkage disequilibrium among loci. We simulated 200 SNP loci in each scenario, and later sampled 96 of them with minor allele frequency *>* 0.1. We recorded all 430 individuals with active infections at the ending time point of simulation in Scenario 1. For Scenario 2 and 3, we randomly sampled 200 infected individuals since the total numbers of infected were too large. The parameters that vary among scenarios are listed in Table S4.

**Table S4.** Parameters for simulated malaria infection data using the stochastic simulator from [32]. The parameters are named according to the literature, where n genes initial means the number of genes in the initial gene pool, and n alleles per locus initial is the initial number of alleles for each epitope locus. Biting rate refers to per day contact rate of a host from vectors, and the actual daily biting rate equals this rate multiplied by daily mosquito population size (we used default values from [32]).

| Scenario | n_genes_initial | n_alleles_per_locus_initial | biting_rate |
| --- | --- | --- | --- |
| 1 | 2000 | 200 | $5 \times 10^{-5}$ |
| 2 | 10000 | 1000 | $1 \times 10^{-4}$ |
| 3 | 20000 | 2000 | $5 \times 10^{-4}$ |

After simulating the infection process, we simulated minor and major allele read counts per host per site to mimic a short-read next-generation sequencing process. On average, the number of total read counts *R*_*ij*_ is around 115, which is typical in high-quality SNP reads datasets. Under the balanced frequency scenario, the number of major allele reads was simulated according to the Poisson model Eq. (3). For the unbalanced strain simulations in Section S1.2, we first sampled the weight of each strain according to a Dirichlet distribution and used the weighted major allele frequency to simulate major read counts, also according to the Poisson model.

In Section 2, before analyzing MOI, we removed the duplicated strains with identical SNP haplotypes. If a host contained multiple strains sharing identical SNP haplotypes, or highly similar SNP haplotypes (for network analysis in figure 1, 2 and S3), they were counted as one strain when calculating MOI. Such removals were not performed in calculating allele frequency and heterozygosity in order not to lose genetic information.

#### S2.4 Preparing Empirical Datasets for SNP-Slice

In Section 2.5, we applied SNP-Slice to SNP categorical data (homozygous or heterozygous) in three regions of Uganda [22, 33]. The filtering of sampled hosts and sites was processed according to the criteria described in [22].

We obtained HIV-1 with-in host dynamics data from [35]. Since SNPs of HIV-1 parasite populations often consist of more than two states, we retained only bi-allelic SNP sites or tri-allelic SNP sites with the least frequent allele less than 1%. The least frequent allele in the retained tri-allelic SNP sites was then filtered and converted to bi-allelic sites. Since each site usually has more than 10,000x depths of sequencing effort, when the proportion of either major or minor allele reads at a site is below 1% of the total reads, we converted that read count to zero to minimize the impact of sequencing errors. Only 3 patients were included in our analysis since the others did not have enough sampling time points in the data. Out of 6,000 sites, we subsampled 96 SNP sites for Patient 1 and 3, and 49 SNP sites for Patient 7 for SS inference.

#### S2.5 Parameter Values for SS and RM in Section 2

##### SNP-Slice

In each scenario, we ran 128 MCMC chains in parallel and all tables and figures include results from these 128 chains. Each MCMC chain continues for a maximum of 10,000 iterations if read count data is used as input (SS-NegBin, SS-Pois, and SS-Bin) and 5,000 if categorical data is used as input (SS-Cat). At least half of the maximum iteration is run as the burn-in process. MCMC is ended when the local maximum likelihood estimator is not changed for 10% of the maximum iteration. The only exception is ‘Nagongera’ of Uganda, where the number of hosts is large and the infection patterns are complicated (462 individuals, most of whom having mixed infections). Due to time constraints, we set up the maximum MCMC iteration to be 500. As seen in Figure 3, using only 500 MCMC iterations of SS-Cat already yields more realistic results than using RM-Cat.

For simulated malaria infection data, since the average MOI values in Scenarios 1-3 are respectively 1.27, 2.58, and 3.90, the concentration parameter *α* is set as 1.5, 3, and 4. However, we have discussed that SNP-Slice is not sensitive to this parameter (Section S1.4). The parameters for the probability of erroneously calling in the categorical method are set to be as low as *e*_1_ = *e*_2_ = 0.001.

For both real infection datasets (HIV and malaria), we use *α* = 3 since SNP-Slice is not sensitive to this parameter. For empirical malaria in Uganda, we follow [22] and use *e*_1_ = *e*_2_ = 0.05.

##### The Real McCOIL

In [22], the total number of MCMC iterations is 100,000 for simulation data. Our trials have found out that setting the total number of MCMC iterations to be 10,000 and 100,000 could generate almost the same summary statistics across 128 chains, so we run 128 chains with 10,000 iterations for all RM cases. The initial MOI is always set to 1.

For empirical malaria infection data in Uganda, we keep using the parameter values used in [22], including *e*_1_ = *e*_2_ = 0.05 and 25 as the upper bound of MOI (maxMOI).

For simulated malaria infection data, we use *e*_1_ = *e*_2_ = 0 when running the categorical method as indicated by [22]. We use maxMOI of 8, 12, and 16 as maxMOI in Scenarios 1-3. The default value of this parameter is 25, but we have shown that RM-Cat is quite sensitive to this parameter (Section S1.5), so here we choose these values to be around or a little less than the maximum inferred MOI by all methods of SNP-Slice to avoid getting extremely high values of MOI from RM.

## Notes

### Competing Interest Statement

The authors have declared no competing interest.

### Summary of Updates

Updated title, and shortened the main text by moving sections to supplemental materials.

